# Patch Testing in Lao medical students

**DOI:** 10.1101/631903

**Authors:** Catriona I. Wootton, Mick Soukavong, Sonexai Kidoikhammouan, Bounthome Samountry, John S.C. English, Mayxay Mayfong

## Abstract

**Background:** Dermatological services in Laos, South East Asia are limited to the capital and patch testing is currently not available, so no data exists regarding the common cutaneous allergens in this population.

**Objectives:** The aim of this study was to document common allergens in medical students in Laos. Patients/Materials/Methods

One hundred and fifty medical students were patch tested using TRUE Test® panels 1 to 3 (35 allergens). Readings were taken at Days 2 and 4.

**Results:** Thirty-eight students (25.3%) had a positive reaction to at least one allergen, accounting for 52 reactions in total. The proportion of the students with positive patch test reading was significantly higher in the female [33/96 (34%)] than in the male [5/54 (9%)], p<0.001. The most common allergens were: nickel (10%), gold (6.6%), thiomersal (6.6%), cobalt dichloride (2%) and p-tert-Butylphenol formaldehyde resin (2%). Balsam of Peru (0.66%), black rubber mix (0.66%), Cl+Me-Isothiazolinone (0.66%), fragrance mix 1 (0.66%), quinolone mix (0.66%), methyldibromo glutaronitrile (0.66%), mercapto mix (0.66%), epoxy resin (0.66%), paraben mix (0.66%), thiuram (0.66%) and wool alcohols (0.66%) accounted for all of the other positive reactions.

**Conclusion:** This study represents the first documented patch test results in Lao medical students and in the adult Lao population. The results of this study will inform any future research into contact allergy in Laos and give an insight into the background level of contact sensitivity in this population.

## Introduction

Laos is a land-locked country in South East Asia with a population of almost 6.8 million, roughly 800,000 of which live in the capital, Vientiane^1^. The country is made up of several different ethnic groups and the main occupation is rice farming. A dermatology clinic exists in the capital but patch testing is currently not available. The aim of this study is to document common allergens in Lao medical students.

## Methods

Year 2 and 3 medical students were selected as the best cohort for the study; it was felt that at this stage of their training they might benefit from learning how to patch test and these year groups are based at the University of Health Sciences, close to the hospital where the patch testing was performed. A lecture on allergy and patch testing was given by the lead author to the Year 2 and 3 medical students, one year group at a time. Following the lecture, the study was explained to the students who were informed about how they could take part. Ethical approval was granted by the Lao National Ethics Committee for Health Research. At the time of testing, the process of patch testing was explained again to each medical student and verbal consent given. A brief questionnaire was completed regarding age, gender, any medical conditions and any known allergies, the TRUE Test® patches (SmartPractice, Denmark: http://smartpractice.dk), 3 panels equalling 35 allergens in total, were then applied to the participant’s upper back. Table 1 lists all of the allergens tested. The participants were reviewed on Day 2 (48 hours) (when the patches were removed) and Day 4 (96 hours). The patch testing was performed and the results interpreted by an experienced dermatologist, following the British Association of Dermatologists guidelines on the management of contact allergy^2^. Those that could not attend the appointments on Day 2 or 4 (5 students in total), were seen within 24 hours or images of their backs were taken for assessment.

**Table 1:**
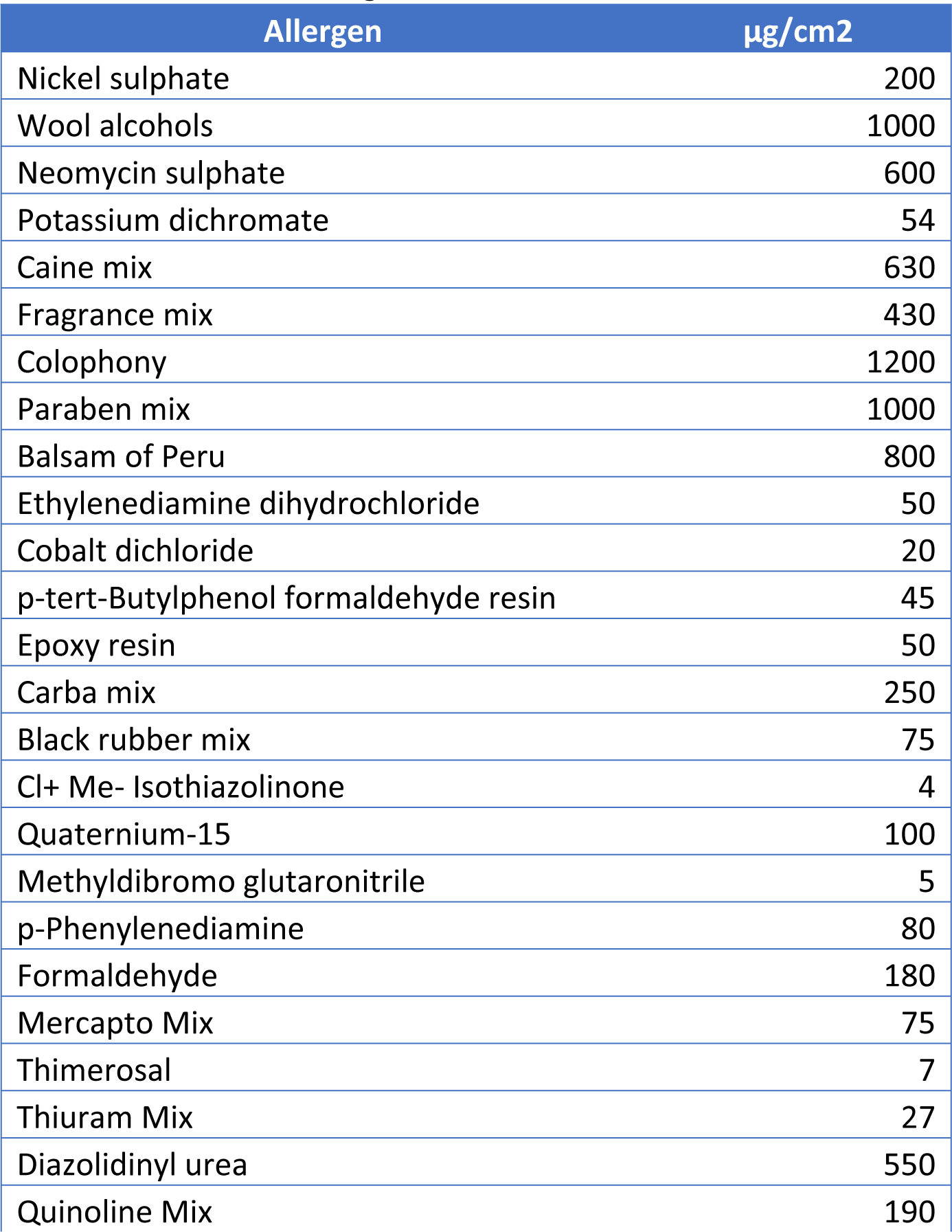
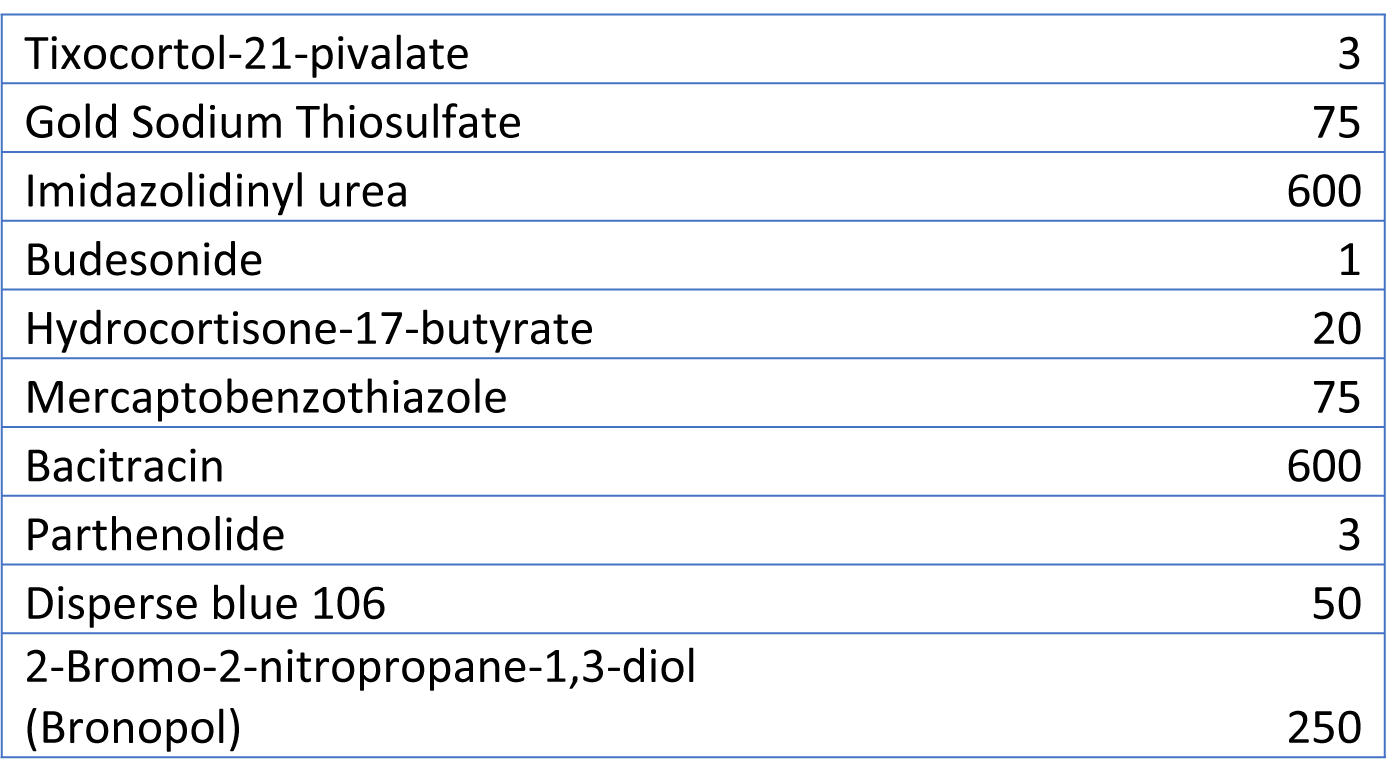
TRUE Test® allergens

All data were recorded anonymously using the Open Data Kit programme on tablet devices. The London School of Hygiene and Tropical Medicine server was used for data storage (http://opendatakit.lshtm.ac.uk).

## Results

One hundred and fifty medical students were patch tested. The age range was from 18 to 35 years, with a median age of 20 years. Sixty-four percent of the participants were female. Ten (6.7%) of participants reported a medical condition: five had asthma, the other conditions included hepatitis B, tinea infection, previous pneumonia and undiagnosed gastrointestinal symptoms. No participant had a diagnosis or evidence of dermatitis. Thirty-nine students (26%) reported a suspected allergy of some form, these included: seafood/shrimp (25.6%), reaction to arthropod bites (17.9%), house dust mite (15.3%), antibiotics or other medications (7.6%), gold (5.1%), jewellery (5.1%), washing powder (5.1%), bee venom, ant venom, frog meat, fermented food, heat rash and alcohol.

Thirty-eight students (25.3%) had a positive reaction to at least one allergen, accounting for 52 reactions in total. The proportion of the students with positive patch test reading was significantly higher in the female [33/96 (34%)] than in the male [5/54 (9%)], p<0.001. The most common allergen was nickel sulphate which resulted in a positive reaction in 15 students (10%). Gold sodium thiosulphate and thiomersal where the next most common allergens resulting in 10 reactions each (6.6%). Cobalt dichloride and p-tert-Butylphenol formaldehyde resin were both positive in 3 students (2%) and Balsam of Peru, black rubber mix, Cl+Me-Isothiazolinone, fragrance mix 1, quinolone mix, methyldibromo glutaronitrile, mercapto mix, epoxy resin, paraben mix, thiuram and wool alcohols all resulted in a single positive reaction (0.66%). Ten students had two reactions (6.6%) and 2 students had 3 reactions (1.3%).

Of the 6 students who reported possible contact allergies (gold, jewellery and washing powder), only 2 had a relevant positive patch test (one to gold and one to nickel).

## Discussion

The point of this study was to patch test medical students in Laos. Our ultimate aim was to try and give an estimate of the background contact sensitivity in the Lao population however generating a cohort which would be truly representative of the adult Lao population and successfully patch testing them would be very challenging and was beyond our capabilities at this time. Medical students were chosen as a suitable cohort as they are generally healthy young adults and were also very engaged in the process as it was a learning experience for them and this helped to ensure a very high follow-up rate.

As our cohort were asymptomatic, the rates of contact sensitivity are lower than in most other documented studies as these typically look at individuals with suspected contact allergy.

The most common allergens were nickel (10%), gold (6.6%) and thiomersal (6.6%). We anticipated that gold and nickel would be common allergens as Boonchai & Iamtharachai^3^ found gold and nickel to be the most common allergens in their review of patch test results over a 9 year period in Thailand. Their rates of positivity were much higher (30.7% for gold and 27.6% for nickel), but their cohort were patients with suspected contact dermatitis. Boonchai & Iamtharachai^3^ highlight the traditional and cultural relevance of gold in Thai society, and gold features in a similar way in the Lao culture with gold jewellery often being used from a very young age and religious items, especially Buddha statues, being covered in gold leaf.

How et al^4^ documented the most common allergens found on patch testing in Kuala Lumpur, their rates of contact sensitivity to nickel, gold and thiomersal were: 35.5%, 15.2% and 14.4% respectively. Again, all of their patients were suspected of having contact dermatitis. The majority of their nickel sensitive patients were female, Boonchai & Iamtharachai^3^ also found that nickel allergy was statistically more common in females; in our study all of the students with reactions to nickel were female and this fits with one of the most common sources of exposure to nickel being costume jewellery^4^. Hamann et al^5^ found that approximately 30% of earrings bought from various locations in China and Thailand were positive for nickel release using the dimethylglyoxime test, confirming the high rate of potential nickel exposure from costume jewellery in this region. Ten (6.6%) of our cohort reported an allergy to eating seafood (over 25% of all those reporting any allergies); positive reactions to nickel have been strongly associated with seafood allergy^6^ this is due to shellfish and some fish containing considerable level of nickel^6^. Despite this only two of the students reporting a seafood allergy had a positive reaction to nickel on patch testing in our study.

Thiomersal was a common allergen in our series, with 6.6% of students having a positive reaction to it. Thiomersal (Merthiolate) is sodium ethylmercurithiosalicylate, an organic mercurial derivative^7^ with antiseptic and antifungal properties, used as a preservative in vaccines, antivenom, ophthalmic and nasal preparations as well as skin prick testing antigens, contact lenses solutions and tattoo ink^8,9^. Contact sensitization to thiomersal is generally regarded as not clinically relevant and related to its use in childhood immunisations, something which has been phased out^7^. As a result, it is expected that positive reactions to thiomersal should decrease in the future and many clinicians no longer include thiomersal in their standard series^7^. However, Yin et al^10^ reported thiomersal as the third most common allergen (11.6%) in their retrospective review of patch testing in Chongqing, China. The authors discuss that positive reactions to thiomersal may be more relevant in their population due to the presence of thiomersal in skin-lightening creams and prohibited tooth fillers and exposure from these products is also likely to be of relevance in the Lao population. Interestingly Moller mentions that the mean age for thiomersal allergy appears to be younger than for many other allergens, being most common around the age of 30 years^7^. Moller also highlights the typical lack of clinical relevance for this allergen with the exception of eye drops, contact lens cleaning solutions and cosmetics, however he does mention a Spanish report where over 50% of the positive reactions to thiomersal were directly related to use of a thiomersal containing disinfectant^7^. Given that our cohort are all medical students (and therefore more exposed to disinfectant and antiseptic preparations) and the relative lack of regulation in products on sale in Laos, it is possible that thiomersal allergy is more clinically relevant in our cohort than elsewhere.

Diepgen et al^11^ performed a cross-sectional study on a random sample of the general population of five different European countries, using TRUE test® panels 1-3 and a few additional allergens from the European baseline series. Many of their findings echo the results from our study. Twenty-seven percent of their cohort had at least one positive reaction; this figure was 25.3% in our cohort. Unsurprisingly, nickel was the most common allergen with a positive incidence rate of 14.5% (10% in our study), thiomersal was the second most common allergen at 5% (6.6%); cobalt 2.2% (2%), p-tert-Butylphenol formaldehyde resin 1.3% (2%) and fragrance mix I 1.8% (0.66%) were also amongst the most common allergens in both studies. Interestingly, no positive reactions to gold were recorded in this cohort, which is in direct contrast to the results of our study where gold was the second most common allergen alongside thiomersal. Although Diepgen et al’s paper^11^ does not specifically mention gold, it is present in the TRUE test® panel 3 and is presumed therefore to have been tested during their study. This paper provides an interesting comparator to our study as they also used the TRUE test® panels 1-3; in our study only 3.9% of the cohort reported a self-diagnosis of possible contact allergy and in Diepgen et al’s study 15.1% of their cohort reported a diagnosis of contact allergy (only 8.2% were physician confirmed)^11^. The most striking difference between the results of these two studies is the incidence of positive reactions to gold in our cohort from Laos; as discussed above, the traditional and cultural use of gold may account for this difference and our findings are in-keeping with others from this region^3,4^.

There are several weaknesses in this study. Firstly, our cohort were all medical students; whilst this ensured our high rate of follow-up it does mean that our cohort is not representative of the Lao population at large. It is worth noting however that the medical students came from all over Laos so a degree of ethic and geographical diversity was present in our sample. Due to the use of medical students, the age range of our cohort was limited to young adults and it would be very interesting to see if our results were replicable in an older Lao cohort. The sister study of this paper looked at contact sensitivity in Lao paediatric patients with atopic dermatitis, giving an indication of common allergens in the younger age group. Secondly, we used all 3 panels of TRUE Test® series, which ensured consistency in dosage but compared to the British Standard Series, the TRUE Test® series does not include p-Chloro-m-cresol, cetearyl alcohol, sodium metabisulfite, fusidic acid, chloroxylenol, *Compositae*, primin, fragrance mix II, kathon CG, methylisothiazolinone, lyral, limonene or linalool, so any sensitivity to these allergens would have been missed. Finally, none of our cohort had evidence of dermatitis so none were suspected of having allergic contact dermatitis, which makes comparison with other studies difficult as the majority of patch test studies consider patients with suspected contact dermatitis. Because of the lack of dermatitis in our cohort, no assessment was made regarding the relevance of any positive patch tests.

## Conclusion

The aim of this study was to document common allergens in Lao medical students. As an asymptomatic cohort of the Lao population, the medical students were chosen in an attempt to give an indication of the background level of contact sensitivity. The medical students proved to be an easy to access cohort who were engaged in the process, ensuring a high follow-up rate. The most common allergens were: nickel (10%), gold (6.6%), thiomersal (6.6%), cobalt dichloride (2%) and p-tert-Butylphenol formaldehyde resin (2%). The most common allergens in this study (nickel, gold and thiomersal) are in-keeping with other findings from this region and it is hoped that the results of this study may help inform future research and the clinical use of patch testing in the Lao population.

## Acknowledgments

This work was funded through the Geoffrey Dowling Fellowship from the British Association of Dermatologists.

